# From Atoms to Fragments: A Coarse Representation for Efficient and Functional Protein Design

**DOI:** 10.1101/2025.03.19.644162

**Authors:** Leonardo V. Castorina, Christopher W. Wood, Kartic Subr

**Affiliations:** AstraZeneca UK, 1 Francis Crick Ave, Cambridge, CB2 0AA, UK; School of Biological Sciences, The University of Edinburgh, Roger Land Building, Edinburgh, EH9 3FF, UK; School of Informatics, The University of Edinburgh, 10 Crichton Street, Newington, Edinburgh, EH8 9AB, UK

**Keywords:** Protein Representation, Protein Design, Protein Search, Fragments

## Abstract

**Motivation:** Although deep learning has accelerated protein design, current protein representations such as sequences or full-atom structures scale non-linearly with protein length. We propose a sparse and interpretable representation for proteins, based on evolutionarily conserved fragments. Specifically, we use a curated set of 40 functional and evolutionarily conserved fragments as an alphabet to build Fragment Graphs and Fragment Sets. These fragment-based representations are both lightweight and functionally informative, capturing up to 55% more variance using fewer than 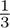 of the dimensions required by traditional methods.

**Results:** On a dataset of 215 functionally diverse proteins, our approach creates more coherent functional clusters than traditional sequence- and structure-based methods, even among proteins with ≤ 30% sequence identity. Fragment-based searches of protein databases achieve accuracies comparable to traditional methods, while using 90% fewer tokens per protein. These searches execute ∼68.7× faster than RMSD-based structural methods and ∼1.64× faster than sequence-based methods, even including fragment pre-processing overhead. Additionally, we show that our representation effectively guides RFDiffusion for protein backbone generation with functional recovery rates higher than 40%. In summary, our fragment-based representation offers a scalable and interpretable alternative for the next generation of protein design tools for backbone design, sequence design, and functional similarity searches within protein structure databases.

**Availability:** https://github.com/wells-wood-research/tessera (Documentation to be made available upon acceptance)

## Introduction

Designing functional proteins could transform medicine, biotechnology, and sustainability. From enzymes that catalyze reactions, to vaccines against target diseases, proteins are precise molecular tools. Yet, designing proteins remains a computationally intractable problem due to the combinatorial nature of the search space, making exhaustive search unfeasible.

Artificial Intelligence (AI) methods enabled *de novo* design of protein binders [12], antibodies [25], and enzymes [30, 20], using diverse representations: language models treat proteins as sequences [10, 23], diffusion models encode backbone geometry as vector frames [28, 2], and other approaches represent structure as voxel grids [7, 21] or graphs [9].

Despite enabling breakthroughs, these representations impose scale non-linearly with protein size, leading to high computational costs and complex models, highlighting the need for efficient and interpretable alternatives.

We propose **fragment-based representations** that model proteins as combinations of evolutionarily conserved structural fragments instead of sequences or full atomic coordinates (See Fig. 1). This approach stems from protein evolution theory, where structures evolved from recombination, repetition, and accretion of small, functional peptides [3]. We show that fragments significantly reduce dimensionality while preserving functional signatures, enabling faster, more interpretable protein search and design.

**Fig. 1.**
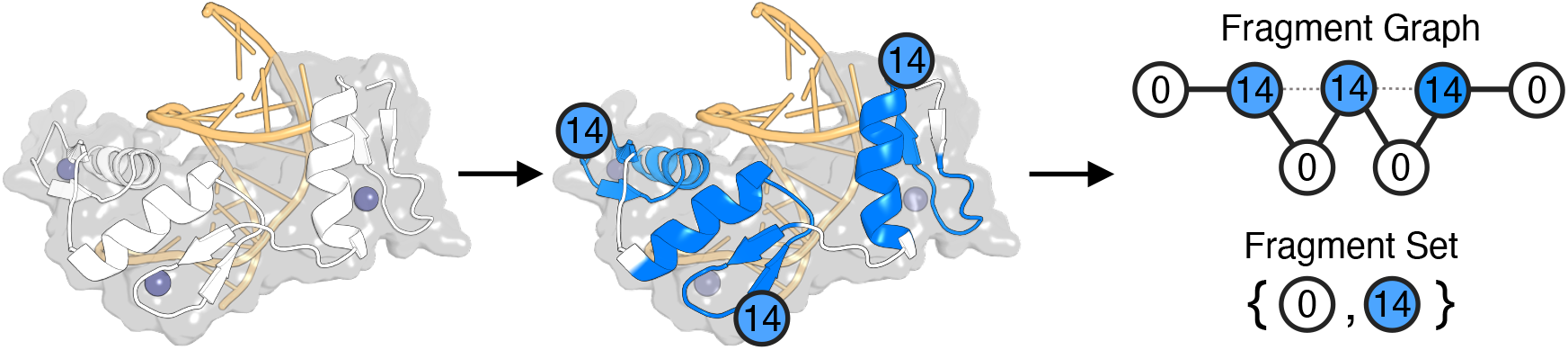
Fragment-based protein representation of the ZIF268 Zinc Finger (PDB: 1AAY). Detected Fragments 14 (in blue), correspond to DNA- and metal-binding functions essential for zinc fingers. Unclassified regions are labeled as “unknown” (white). The structure is represented as a Fragment Graph, which preserves connectivity via peptide bonds (dark edges) and spatial proximity (dotted edges) or as a Fragment Set, with unique fragment types. DNA shown in yellow; zinc ions in purple.

Proteins naturally lend themselves to abstraction at multiple scales, from secondary structures such as *α*-helices and *β*-sheets [22] to tertiary structural motifs strongly associated with molecular functions such as *β* hairpins [19]. Our fragment-based representation continues this trend, using fragments as functional building blocks rather than full atomic structures.

Previous studies have identified and leveraged recurring structural motifs for protein analysis and design. Alva et al. [4] identified conserved ancient fragments associated with binding functions. Frappier et al. [11] introduced recurring Tertiary Structural Motifs (TERMs) to design protein binders. Kolodny et al. [17] compiled several sequence-based “THEMES” for functional protein analysis.

Building on these ideas, our method explicitly encodes the backbone geometry rather than sequences or atomic coordinates, decomposing proteins into functional fragments. While our method is library-agnostic and supports custom structural motifs, here, we validate our approach using the minimal alphabet of 40 ancient fragments described by Alva et al. [4]. We selected these fragments because they are evolutionarily conserved in modern functional families, such as DNA- or Metal-binding proteins, making them ideal candidates for abstracting complexity while retaining functional meaning. We demonstrate three key applications of our fragment-based approach: (1) functional clustering to evaluate how well fragments capture protein function; (2) database searching to demonstrate effectiveness in retrieving functional proteins and computational efficiency; and (3) protein design using fragments as blueprints to guide RFdiffusion to generate backbones with functional signatures.

We also provide a vectorized Python package (MIT License) for fragment representations, adaptable to any fragment library.

## Methods

We propose to represent proteins abstractly, as compositions of fragments. We chose the 40 ancient functional fragments identified by Alva et al. [4], because they are a curated and ubiquitous vocabulary of peptides that are evolutionarily conserved and explicitly linked to protein function. Here, we first explain fragment-based representations and then evaluate them via functional clustering, database searching, and protein design, comparing against traditional representations.

### Fragments as a Coarse Representation of Proteins

Given a protein structure from a PDB file, our representation decomposes it using evolutionarily conserved fragment motifs. We propose two representations for the protein structure, without sequence information, as a **Fragment Graph** or as a **Fragment Set**. In the former, nodes in the graph represent fragments and edges denote peptide bonds or spatial proximity. Fragment Sets, on the other hand, contain lists of unique fragments present, regardless of their arrangement (See Fig. 1).

To construct the reference library, we extracted all instances of the 40 fragments from Alva et al. [4] from PDB structures using AMPAL [29]. Then, we filtered out fragments with erroneous labeling or lengths differing from the canonical definitions, yielding 219 verified instances (See Supp. Table 1). These are used as “ground truth” by our detection algorithm for all subsequent search, clustering, and design experiments.

### Building Fragment Representations

We build fragment representations by: (1) detecting the fragments in the structure, (2) classifying unmatched regions, and (3) converting to a graph or a set (See Fig. 1).

Fragment detection identifies segments of the target protein matching library fragments below a distance threshold using a sliding window algorithm (see Supp. Section 1 and 2). For this distance calculation, we evaluated several distance metrics:

- **Sequence-based** (sequence identity, BLOSUM distance) measure distance in amino acid sequence [14].
- **Angle-based** (RMS, RamRMSD, LogPr) are sequence-independent and measure distance in backbone torsion angles (*ϕ* and *ψ*) [16].

This yields a normalized **Fragment Distance Matrix** *D*, where each entry *D*_*ik*_ denotes the distance between index *T*_*i*_ of the target structure and fragment *F*_*k*_ from the library. A segment is classified as matching a fragment if *D*_*ik*_ *<* 3.65% (See details in Supp. Section 5). If fragment matches overlap, we retain the one with lower distances, allowing up to two amino acids overlap between adjacent fragments.

Unmatched regions are classified as **unknown fragments** (9–24 residues) or **unknown connectors** (*<* 9 residues). Finally, we represent classified regions as:

- **Fragment Sets:** presence or absence of fragment classes without considering connectivity information.
- **Fragment Graphs:** graphs with nodes for each fragments and edges for peptide bonds or spatial proximity (*<*10 Å). Edge features are one-hot encoded by types.

Fragment Sets are suited for speed-critical tasks like search, while Fragment Graphs preserve structural context, useful for clustering and design.

### Datasets for Validation

To validate whether fragments preserve structural and chemical properties, we used **PDBench** [6], a fold-balanced structures dataset.We quantified properties such as hydrogen bonding and solvent accessibility in fragment versus non-fragment regions, normalizing for fragment density to compute enrichment ratios (detailed calculation methods are provided in Supp. Section 9). For each property, we computed the correlation between its proportion in fragment regions and the overall fraction of the protein covered by fragments.

To evaluate functional clustering, database search, and design, we curated the **Protein Function Dataset (PFD)**, including 215 protein monomers across 12 functional binding classes (DNA, RNA, metal ions, GTP, ATP, and combinations). Structures were filtered using Gene Ontology (GO) codes and ≤ 30% sequence identity via the PDB Advanced Search [5]. Where possible, we selected 10 representative structures per functional category (detailed in Supp. Table 2). We use this to evaluate functional clustering, database search, and design.

### Fragments for Functional Clustering

We evaluated whether fragment-based representations better capture functional similarities between proteins compared to traditional structure- and sequence-based approaches. Specifically, we assessed whether proteins with similar functions form more coherent clusters under different representations. For all protein pairs in the PFD, we computed pairwise distances using four metrics:

1. **RMSD (Root Mean Square Deviation)**: Structural distance using BioPython’s CEAligner [26, 8].
2. **BLOSUM62**: Sequence distance using pairwise alignment scores based on amino acid substitution frequencies [14].
3. **BagOfNodes**: Set distance based on the presence or absence of fragments (Frag. Sets).
4. **GraphEditDistance**: Graph distance accounting for fragment identity and spatial arrangement (Frag. Graphs).

Then, to evaluate the information density of each representation and assess its potential for compact vector-based indexing, we projected each protein into a distance-preserving latent space of increasing dimensionality using Principal Coordinate Analysis (PCoA) and t-SNE.

To quantify clustering performance, we applied Gaussian Mixture Models (GMM) and K-Means with 12 clusters (matching known functional categories in PFD)on 2D projections of the embeddings. To measure robustness and to evaluate clustering quality, we use Adjusted Rand Index (ARI) and Normalized Mutual Information (NMI) to compare against ground truth labels. To assess the intrinsic quality of the embeddings, we calculated Silhouette and Trustworthiness scores, F1-score, and the correlation coefficients using the respective native distance matrices for each representation (e.g., RMSD for structure, BLOSUM for sequence).

### Fragments for Functional-based Searches

While the previous experiment evaluated the quality of the representation, here we assessed whether fragment-based representations support fast and meaningful protein database searches. Specifically, we measured retrieval speed, memory requirements, and functional relevance of the results.

We benchmarked fragment-based methods GraphEditDistance and BagOfNodes, against traditional methods RMSD and BLOSUM) querying 1, 10, and 100 proteins from a database of 100 structures using 35 cores ^1^. For each method, we recorded initialization time, query time, and memory usage, measured as the number of data points per protein.

To assess retrieval quality, we queried each PFD protein against all others and ranked the results by distance. We evaluated whether functionally similar proteins ranked higher using Normalized Discounted Cumulative Gain (NDCG) and Area Under the Receiver Operating Characteristic (AUROC) (see Supp. Section 8).

### From Fragments to Functional Proteins

We explored using fragments as blueprints for generating functional proteins by providing structural constraints to RFDiffusion. We hypothesize that if fragments capture functional information, then fragment-derived templates should yield structures similar to proteins with the same function.

To test this, we used RFDiffusion[28], a state-of-the-art backbone generation model to complete partial templates of the 215 PFD proteins, where fragment backbone coordinates were held fixed, while intervening non-fragment regions were removed. RFDiffusion then regenerated the missing regions, producing five candidate backbones per protein. We used each generated structure as a query in FoldSeek to retrieve the top 10 structurally similar proteins from the PDB using sequence-independent search [27].

To assess functional recovery rate, we calculated the the fraction of generated designs whose top 10 matches shared the exact Gene Ontology (GO) code(s) of the original protein. As a baseline, we also evaluated with unmodified backbones.

To determine if functional recovery was driven by evolutionary signals or just geometric constraints, we used a naïve control using 40 random segments extracted from the same proteins as the fragments in [4]. These fragments matched the length distribution of the evolutionary library and were used to guide generation under the conditions described earlier.

## Results

We evaluated our fragment-based representation across four key areas: fragment detection accuracy, physicochemical properties of fragments, effectiveness in capturing functional patterns, and applications in protein search and design.

We validated our fragment detection algorithm using both sequence-based (BLOSUM and Sequence Identity) and angle-based (LogPr, RamRMSD, and RMSD) distance metrics. RMSD, via both PyMol and BioPython, was significantly slower and prone to silent failures during alignment.

As shown in Supp. Fig. 1, individual metrics achieved modest F1 scores (∼0.40). However, combining two complementary metrics, such as LogPr and RamRMSD, significantly improved performance to ∼0.85. Adding a third metric provided no significant improvement.

Using Receiver Operating Characteristic (ROC) analysis, we identified an optimal distance threshold of 3.65% for fragment classification, achieving an AUROC of 87% (See Supp. Fig. 2).

### Fragments Show Distinct Structural and Chemical Properties

We compared fragment and non-fragment regions across the PDBench benchmark to determine whether fragments capture distinct structural or chemical properties.

As shown in Supp. Fig. 6, fragments covered ∼40% of each protein on average with consistent standard deviations. There were some outliers like Alpha Solenoid or Alpha-Beta Horseshoe with lower coverage (∼20%) and the special folds had the highest standard deviation. Fragment coverage was consistent across resolutions (Supp. Fig. 7).

Fragment regions had a higher proportion of intra-fragment hydrogen bonds, especially in mainly *β* folds with a ∼15% increase compared to non fragment regions. In contrast, inter-fragment hydrogen bonds were lower, particularly in the mainly *α* folds with a ∼ 47% decrease. Surface accessibility was also slightly reduced in fragment regions (∼5%) across most folds except special folds. Charge, polarity, and secondary structure distribution remained comparable between fragments and non-fragment regions (see Supp. Section 9).

### Fragment-Based Embeddings Efficiently Capture Functional Similarities

We evaluated how well fragment-based representations and traditional sequence-and shape-based methods capture functional similarities in reduced-dimensional embeddings. We compute a distance matrix for the PFD using RMSD (shape), BLOSUM (sequence), BagOfNodes (fragment sets), and GraphEditDistance (fragment graphs).

We projected the distances into distance-preserving latent-spaces using PCoA and calculated the cumulative explained variance across dimensions (Fig. 2A). Fragment-based embeddings preserved significantly more cumulative variance than traditional metrics, with BagOfNodes and GraphEditDistance preserving over 95% and 80% within 20 dimensions, respectively. In contrast, BLOSUM and RMSD preserved less than 60% and 40%, respectively.

**Fig. 2.**
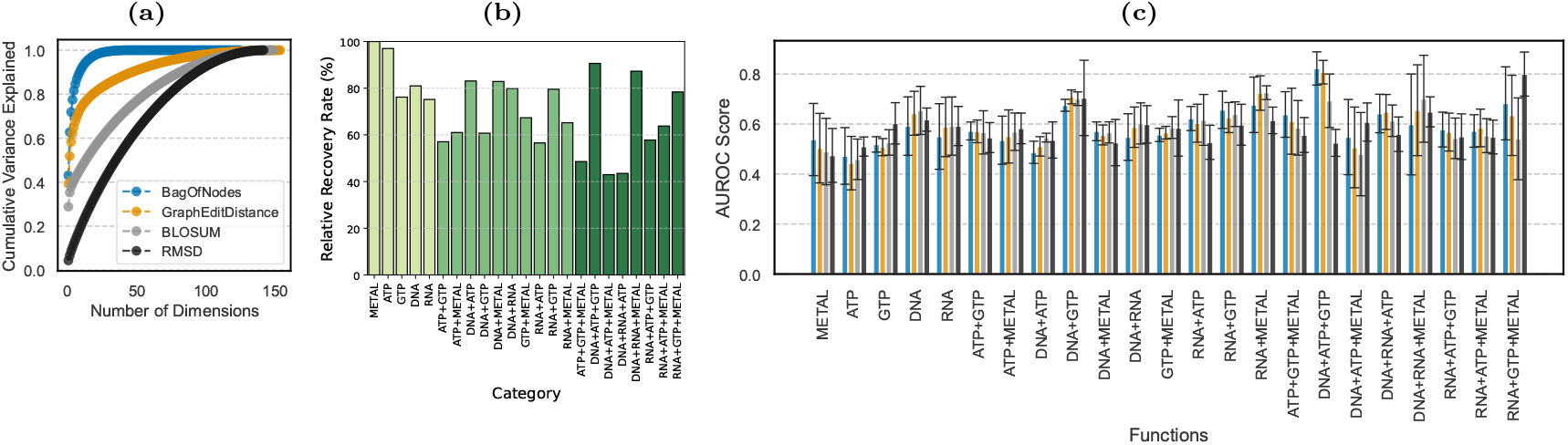
Quantitative evaluation of fragment-based representations. **(a)** Cumulative variance explained by different distance metrics after Principal Coordinate Analysis (PCoA) projection of the Protein Function Dataset (PFD) containing 215 proteins across 12 functional categories. **(b)** Relative functional recovery rates for fragment-guided backbone generation. We used fragments as fixed constraints for RFDiffusion. We queried the generated backbones against the PDB using Foldseek with TM-Align mode to compare the functional annotations of the top structural matches against the original backbones. **(c)** Area Under the Receiver Operating Characteristic Curve (AUROC) scores for protein similarity search across different distance metrics. Higher values indicate better separation between relevant and non-relevant search results.

Despite being sequence-agnostic, fragment-based distances correlated well with sequence-based distances. GraphEditDistance achieved a Spearman correlation of 0.91 with BLOSUM, and BagOfNodes 0.57. RMSD, however, showed weak or slightly negative correlation with other metrics.

We evaluated clustering performance using GMMs and K-Means on PCoA, t-SNE, and UMAP embeddings (See Table 2 and Supp. Section 7). Overall, fragment-based representations demonstrated better clustering performance across most metrics.

Notably, BagOfNodes achieved the highest Silhouette score (0.8227), indicating well-separated clusters, along with the best Silhouette and Trustworthiness scores, and Distance Correlation. GraphEditDistance had the highest ARI (0.0458) and F1 score (0.1985), indicating the highest agreement with the true functional clusters after adjusting for chance. RMSD ranked highest in NMI score (0.3863), reflecting better mutual information between cluster assignments and true functions, and was second best for ARI and F1 Score. BLOSUM ranked second in Silhouette scores and Trustworthiness scores (See Table 2).

### Fragment-Based Search: Fast and Accurate

We tested functional search retrieval quality and computational efficiency for fragment distance methods (GraphEditDistance and BagOfNodes) against traditional sequence (BLOSUM) and shape (RMSD) distance methods.

We selected each of the 215 annotated proteins from PFD as queries, computing the pairwise distance to all other proteins. We assessed retrieval quality using NDCG and AUROC to measure how highly functionally similar proteins ranked.

As shown in Figure 2C and Supp. Fig 4, all methods performed similarly overall, with scores across within one standard deviation. In terms of retrieval accuracy, fragment-based methods matched approaches for most functional categories, with particularly strong AUROC performance in identifying DNA+ATP+GTP-binding proteins (values *>*0.8 against 0.75 0.56 for RMSD and BLOSUM). RMSD showed an advantage in NDCG for specific functions, especially in DNA+GTP, RNA+GTP, and RNA+GTP+Metal binding searches.

Then, we benchmarked the computational efficiency of each method, measuring query times for 1, 10, and 100 queries against a database of 100 proteins using 35 cores (Table 1).

**Table 1.**
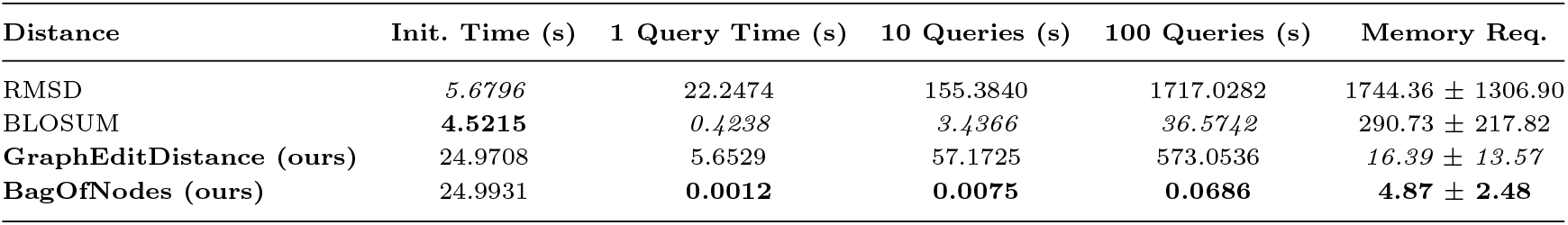
Performance comparison of query methods across different protein representations using the PFD Dataset. Initialization and query times (1, 10, and 100 queries) are measured on a database of 100 proteins using 35 cores. Memory requirement is reported as the average number of data points required to represent a protein: backbone atoms (RMSD), residues (BLOSUM), nodes (Fragment Graphs), or elements (Fragment Sets)(see distributions in Supp. Fig. S26).

**Table 2.** Clustering performance comparison of different distance metrics using Gaussian Mixture Models (GMM) on Principal Coordinate Analysis (PCoA) embeddings of the PFD (215 proteins across 12 functional categories).

| Distance | Based On | ARI | NMI | Silhouette | Trustworthiness | Distance Corr. | F1 Score |
| --- | --- | --- | --- | --- | --- | --- | --- |
| RMSD | Shape | 0.0357 | **0.3863** | -0.0334 | 0.9598 | 0.8647 | 0.1957 |
| BLOSUM | Sequence | 0.0027 | 0.2933 | 0.0248 | 0.9923 | 0.9966 | 0.1640 |
| **GraphEditDistance (ours)** | Fragment Graph | **0.0458** | 0.3832 | 0.0766 | 0.9915 | 0.9829 | **0.1985** |
| **BagOfNodes (ours)** | Fragment Set | 0.0050 | 0.3455 | **0.8227** | **0.9991** | **0.9998** | 0.1660 |

Fragment-based representations use significantly fewer data points compared to traditional methods. Relative to backbone atom-based representations (RMSD), fragment graphs and fragment sets reduce dimensionality by approximately 99.1% and 99.7%, respectively. Compared to sequence-based representations, the reductions are 94.4% and 98.3%, respectively (See Table 1).

Overall, sequence search with BLOSUM distance is the fastest method considering initialization time and search time. BagOfNodes is the fastest search method overall, completing 100 queries in under 0.07 s, while other methods required substantially longer – RMSD took about 1717 s, GraphEditDistance about 573 s, and BLOSUM took 36.57 s.

Although fragment-based methods have a higher initial cost which involves converting protein structures to fragment graphs (around 6 s compared to 25 s), this cost is quickly offset by the faster search times.

### Functional Design Recovery with Fragment-Based Diffusion

We evaluated the ability of fragment-based templates to guide the generation of functional proteins using RFDiffusion. For each of the 215 proteins, we generated a template backbone of fragments. We used RFDiffusion to fill in the gaps between the fragments and generate 5 different structures. Then, we use FoldSeek to search for the closest 10 backbones using sequence-independent TM-Align. For each design and for the original backbone, we define recovery rate as the fraction of backbones annotated with the function of the original backbone (See Supp. Section 10). We also calculate the relative recovery rate as the recovery rate of the design over the recovery rate of the original backbone(See Figure 2B and Supp. Section 10).

In general, there is a range of recovery rates, across different functional categories. Metal-binding proteins achieved perfect recovery rates, while ATP- and GTP-binding proteins also showed consistently high recovery rates. Multi-functional proteins demonstrated more variable outcomes, with DNA+ATP+GTP-binding showing the widest range, varying from 0% to 300% relative recovery rate and also the lowest recovery rate for the control.

Single-function designs generally demonstrated higher recovery rates compared to their multi-functional counterparts. Among dual-function proteins, metal-binding combinations proved most successful, with DNA+Metal, GTP+Metal, and RNA+Metal showing particularly high recovery rates. Interestingly, some triple-function combinations achieved surprisingly high recovery rates, particularly for DNA+ATP+GTP, DNA+RNA+Metal, and RNA+GTP+Metal binding.

To control for the impact of library choice, we also repeated the experiment using 40 naive fragments, generated randomly from the same proteins as our original library. As shown in Supp. Section 10, while Alva et al.[4] fragments recovered proteins across all functional categories, the naive fragments recovered only a few.

### Fragments and Functional Similarity

We identified two DNA-binding proteins with high sequence and shape distances but low fragment distance (Supp Fig. 27). These proteins are the UvrABC system protein C, involved in DNA repair (PDB: 2NRR), and a viral DNA-dependent RNA polymerase (PDB: 6RIE). Despite their overall differences, they share fragments 17 (metal-binding), 23 (nucleotide-binding), and 35 (structural).

Fragment 17 is a small helix involved in metal binding, while fragment 23 is a helix-loop-sheet associated with nucleotide binding. Fragment 35, a sheet-loop-sheet motif, contributes to structural integrity. Notably, none of these fragments are explicitly classified as DNA-binding, yet their presence captures similarities in overall fold architecture. This is reflected in their low fragment distance scores compared to sequence (90% divergence) and shape (RMSD: 6.71 Å) distances.

Additionally, Graph Edit Distance (GED: 4%) accounts for the fragment neighborhood, considering factors such as the number of adjacent unknown fragments and peptide bonds.

## Discussion

In this study, we demonstrate that fragment-based representations effectively coarsen protein structures while preserving essential functional information. Using just 40 evolutionarily conserved fragments, our approach captures important structural properties while reducing the dimensionality by up to 99% compared to traditional methods. Our fast and vectorized fragment-detection algorithm allows fast conversion to fragments and achieves an F1 score of 0.85. Furthermore, we successfully use fragments to guide backbone towards preserving protein functional signatures with recovery rates between 40-100%. These results make fragment-based representations a promising alternative to traditional sequence- and structure-based approaches for protein analysis, search, and design.

### From Libraries to Detection: Building Fast and Robust Fragment Algorithms

We evaluated several metrics for fragment detection, focusing on shape and sequence. While individual metrics performed similarly, combining LogPr and RamRMSD nearly doubled the F1 score from 0.40 to 0.85 [16]. This highlights that torsion-angle alone can outperform sequence-based methods for robust fragment detection.

While both metrics measure differences in backbone torsion angles *ϕ* and *ψ*, they process information differently. RamRMSD uses root-mean-square deviation, where squared differences are averaged, giving more weight to larger deviations. In contrast, LogPr applies a logarithmic transformation to normalized angle differences, emphasizing small deviations and converting them to a probability-like scale. This complementarity allows our algorithm to be sensitive to both, large and small deviations in torsion angles.

The fragment detection algorithm is written in Python and uses AMPAL [29] for parsing protein structures and NumPy’s vectorised convolutional operations. We deliberately avoided structural alignment methods based on incremental combinatorial extension (CE), which, despite potentially improving detection accuracy, proved computationally expensive and occasionally unstable during testing. [26, 8].

A key strength of our software is its flexibility. Users can easily swap our library with custom fragments by providing folders of PDB structures with the fragments of interest. This extends the software applications beyond the binding functions presented here, including enzyme design, antibody engineering, and *de novo* structural design. The software is written in Python and it is highly modular, meaning that users can expand it to integrate their own distance algorithms. Additionally, we use vectorized operations through NumPy, delivering fast performance while retaining the intuitive syntax that Python offers.

### Fragment-based Representations Capture Functional Information

Using the fold-balanced PDBench dataset, we found that fragment regions capture distinct structural and chemical properties. These regions contained higher proportions of intra-fragment hydrogen bond, particularly in mainly *β* structures (+15%). This is consistent with *β* folds forming hydrogen bonds between adjacent strands[24]. On the other hand, inter-fragment hydrogen bonds were significantly lower in fragment regions, with a 47% reduction in mainly *α* structures. This observation is consistent with the characteristic hydrogen bonding pattern of *α*-helices, where hydrogen bonds stabilise the helical structure internally (*i, i* + 4 pattern), reducing the potential for hydrogen bonds with adjacent fragments [24]. These results suggest that fragments may capture “self-contained” structural units. This is also supported by the reduced surface accessibility in fragment regions, such as the core of the protein, which is more likely to have folded regions, compared to surface exposed areas like loops [24].

Additionally, fragment-based representations outperform or match traditional methods in capturing functional similarities in embedding spaces. Both Fragment Graph (GraphEditDistance) and Set (BagOfNodes) metrics consistently achieved strong clustering scores, with BagOfNodes reaching a Silhouette score of 0.82 and GraphEditDistance showing the best overall performance for ARI (0.046) and F1 score (0.20). Fragment-based methods also preserved substantially more information at lower dimensions, achieving 95% and 80% cumulative variance compared to 60% and 40% for traditional sequence- and shape-based methods, respectively at 20 dimensions.

Notably, fragment-based representations capture these functional patterns without relying on the amino acid sequences. Instead they rely solely on backbone geometry. This effectiveness arises because fragments capture functional motifs regardless of their sequential arrangement, which is typical of other alignment-free analysis tools [31]. Sequence-alignment tools assume colinearity, meaning that they expect homologous residues to occur in the same order in both sequences [31]. Structural-alignment tools, such as combinatorial extension (CE), mitigate this by breaking the structure into smaller regions and reassemble them to complete the alignment [26]. However, these tools may struggle when there is little structural homology between proteins with the same function [13]. This may explain why RMSD was poorly correlated with all the other metrics, since the PFD is filtered for ≤ 30% sequence identity, decreasing the effectiveness of a global structural alignment. In contrast, fragment-based representations maintain performance by focusing on the presence of specific functional units. For example, Fragment Sets simply track the presence or absence of functional fragments, without their spatial precise arrangement.

### Fragments for Searching Protein Databases

Protein database searches are essential tools for finding structurally or functionally similar proteins. Traditional sequence- and shape-based methods can miss important relationships when functional motifs are arranged differently, for example when divided by other structural elements. Fragment-based representations overcome this limitation while delivering equal or better performance than traditional methods. Fragments require an initial processing cost to convert structures to graphs or sets. However, this one-time computation can be done for the entire dataset in advance and it is quickly offset by the search speedups. In our benchmarks, Fragment Sets searches using BagOfNodes execute at fractions of a second (0.07s for 100 queries), over 500*×* faster than sequence-based BLOSUM searches (36.57s). Similarly, Fragment Graphs searches with GraphEditDistance are ∼3*×* faster that structure-based searches with RMSD (573s vs. 1717s for 100 queries), but are slower than sequence searches.

The major advantage, however, is memory efficiency. Fragment representations reduce the memory requirements by 90-99%, compared to traditional representations. For large-scale applications involving millions of proteins, this reduction could enable searches on hardware that would otherwise be insufficient for atom- or residue-level comparisons. These efficiency gains, coupled with comparable functional retrieval accuracy (as measured by AUROC and NDCG scores), make fragments an attractive alternative for the next-generation of protein search tools.

### Fragment Constraints as Design Guides

The current generative tools for protein design are difficult to prompt for functional generation. For example, Ingraham et al. [15] highlight that there is currently no protein design system that can: (1) sample conditionally under diverse design constraints without retraining for new target functions, (2) with a sub-quadratic scaling computational efficiency, and (3) which integrates both sequence and structure modeling. For instance, RFDiffusion, a state-of-the-art diffusion model, lacks explicit mechanisms to enforce specific functional constraints in the generated structures.

Fragment-based constraints address this limitation by using evolutionarily conserved functional units to guide the generation process. Instead of retraining models with additional functional labels, our approach leverages evolutionarily conserved “building blocks” to steer generative models toward functionally relevant backbones.

We successfully generated functional-looking protein structures using fragments as RFDiffusion templates. Using our fragment-detection algorithm, we detected fragments in existing proteins and created partial backbones containing only these regions.

We then used RFdiffusion to fill the connecting segments and used FoldSeek to retrieve the closest proteins available. On a dataset of various functional categories, we successfully generated structures that maintained the functional signatures of the original proteins. Recovery rates varied by functional category, with metal-binding and ATP-binding proteins achieving nearly perfect recovery (∼100%). Our approach was particularly effective for certain multi-functional proteins, with DNA+ATP+GTP and DNA+RNA+Metal combinations showing surprisingly high recovery rates despite their complexity. Crucially, when we repeated this experiment using randomly selected “naive” fragments, most functions had 0% recovery rates. This implies that functional recovery is driven by specific, evolutionarily conserved structural motifs rather than random geometric constraints.

These results suggest that diffusion models have implicitly learned about evolutionarily conserved fragments and are able to use them for design. Explicitly incorporating fragment representations in these models could help reduce the computational complexity while also providing more direct functional control to generate specific functional proteins.

### Limitations and Future Work

Our current implementation uses a curated library of 40 fragments spanning functions of DNA, RNA, GTP, ATP, and Metal binding. Consequently, the method’s generalization to other protein classes depends on the fragment repertoire used. For instance, domains such as antibodies require specific fragment sets to capture their hypervariable loop diversity, as shown by recent fragment-based design approaches [1]. Further studies could explore generalization as a function of different fragment libraries or data-driven approaches to discover novel fragments with unsupervised learning, expanding the representation beyond the functions explored here.

Additionally, the high variance captured by our low-dimensional projections (Figure 2), coupled with the significant speedups and lower memory requirements (Table 1) suggests that fragments are well-suited for continuous vector embeddings. Future work could leverage this for ultra-fast vector search, akin to the structural alphabets used by FoldSeek[27].

A major advantage of our approach is its inherent interpretability. Unlike traditional black-box methods such as sequence embeddings[23], fragment-based representations provide clear functional insights as they are associated with specific structural motifs and known biological roles. This interpretability could improve generative models by making their outputs more functionally interpretable and allow more control during the design process. Additionally, for protein sequence design, predicting sequences for entire fragments rather than individual amino acids could improve computational efficiency and better capture the specific sequence constraints associated with each functional motif.

While the BagOfNodes approach is very fast, it is less effective when multiple instances of a fragment contribute to distinct functional roles. For example, Zinc finger proteins usually contain three instances of fragment 14, each binding a positively-charged Zinc ion, and all binding negatively-charged DNA (See Fig. 1). More instances of fragment 14 may indicate binding to multiple DNA strands or different regions of the same strand. In these cases, Fragment Graphs with GraphEditDistance provide a more nuanced representation by capturing the connectivity and fragment context, despite their higher computational cost.

Our fragment detection algorithm achieves a good F1 score of 0.85, but is potentially sensitive to subtle variations in torsion angles which could lead to misclassification. For example, a large change in torsion angles of the middle amino acid of a fragment would change the backbone angles for one amino acid only, so it might still be classified similarly. Future work could integrate probabilistic models to quantify detection confidence and providing adjustable sensitivity, allowing users to choose the settings based on their design scenario.

Finally, our current representation is static, capturing the structural potential for function rather than its dynamics. Protein function often relies on conformational changes and mobility between domains. Recent work suggests that reused structural segments are indeed modular units of protein dynamics [18]. Future work could explore “dynamic fragment graphs” that encode movement and relative flexibility, providing a more complete functional representation.

## Conclusion

We introduced a fragment-based protein representation that encodes structures using a curated library of 40 evolutionarily conserved functional fragments. This approach reduces dimensionality by up to 99% while preserving functional and structural information. Our evaluations demonstrate that fragment-based representations capture functional relationships more effectively than traditional methods in clustering, enable significantly faster database searches with comparable accuracy, and successfully guide RFDiffusion to generate backbones with functional signatures. Unlike black-box representations, our method provides interpretability by linking fragments to biological functions. Fragment-based representations offer a scalable and biologically relevant framework for protein design. By balancing efficiency and interpretability, this approach lays the foundation for the next generation of protein design tools.

## Footnotes

1 Intel(R) Core(TM) i9-10980XE CPU @ 3.00GHz

