## Supplementary Materials for "From Atoms to Fragments: A Coarse Representation for Efficient and Functional Protein Design"

#### 1 Sliding window algorithm

Fragment detection using a sliding window approach. The algorithm scans the target protein structure $T$ using a fragment $F_k$ of length $n_k$, computing a similarity score at each valid position using a specified distance metric (e.g., LogPr for torsion angles or sequence-based metrics). The step size $s$ controls the stride of the window. The output is a Fragment Distance Matrix $D$, which stores similarity values for each alignment position.

**Algorithm 1** Fragment detection via sliding window

**Require:**
  Target structure $T$ of length $L_T$
  Fragment $F_k$ of length $n_k$
  Distance Metric function Metric$(\cdot, \cdot)$ (e.g., LogPr)
  Step size $s$ (default: $s = 1$)
**Ensure:**
  Fragment Distance Matrix $D$ containing similarity values for each valid alignment

```
Initialize D as an empty list
for i ← 1 to L_T − n_k + 1 in steps of s do        ▷ Slide fragment F_k across T
    Extract data from T[i : i + n_k − 1] (angles or sequence)
    Compute similarity: D[i] = Metric(T[i : i + n_k − 1], F_k)
end for
return D
```

#### 2 Fragment Detection Metrics and Details

For computational efficiency and robustness, we avoided structural alignment methods based on CEAligner like BioPython [3]. These methods were computationally expensive and, in our tests, exhibited instability and silent failures when applied to large-scale fragment detection.

##### 2.1 Sequence-based distances

###### Sequence Identity

Sequence identity quantifies the fraction of residues that match exactly between a reference sequence and a fragment sequence. The metric is computed as the proportion of identical residues across all positions, where a value of 1 indicates a perfect match, and 0 indicates no matches.

$$\text{Sequence Identity} = 1 - \frac{\sum_{k=1}^{n} \text{match}(s_k^{\text{ref}}, s_k^{\text{frag}})}{n},$$

where:

- $n$: Total number of residues being compared.
- $s_k^{\text{ref}}$: Residue at position $k$ in the reference sequence.
- $s_k^{\text{frag}}$: Residue at position $k$ in the fragment sequence.
- $\text{match}(x, y)$: A function that returns 1 if the residues $x$ and $y$ are identical and 0 otherwise.

**BLOcks SUbstitution Matrix (BLOSUM):**

The BLOSUM metric measures the similarity between two sequences using the BLOSUM62 substitution matrix. Each residue pair is scored based on its substitution score in the matrix. A positive score indicates a likely substitution, while a negative score indicates an unlikely substitution [4].

$$\text{BLOSUM Distance} = 1 - \frac{\sum_{k=1}^{n} \text{BLOSUM62}(s_k^{\text{ref}}, s_k^{\text{frag}} > 0)}{n},$$

where:

- $n$: Total number of residues being compared.
- $s_k^{\text{ref}}$: Residue at position $k$ in the reference sequence.
- $s_k^{\text{frag}}$: Residue at position $k$ in the fragment sequence.
- $\text{BLOSUM62}(x, y)$: Lookup function in the BLOSUM62 matrix, returning the substitution score for residues $x$ and $y$.

##### 2.2 Angle-based distances

**Root Mean Square (RMS)**

The RMS metric measures the delta in root mean square difference between the fragment's angles and the corresponding angles in the target structure. Smaller values indicate a closer match.

$$\text{RMS} = \sqrt{\left(\Delta\phi_j^2 + \Delta\psi_j^2\right)}$$

Where:

- $\Delta\phi_j^2$: Squared difference of $\phi$ angles between the fragment and target at position $j$.
- $\Delta\psi_j^2$: Squared difference of $\psi$ angles between the fragment and target at position $j$.

**Ramachandran RMSD (RamRMSD):**

The RamRMSD is an aggregate metric that accounts for the RMS distances of the $\phi$ and $\psi$ angles across all $n$ residues of interest, derived from their positions on the Ramachandran space [5].

$$\text{RamRMSD} = \sqrt{\frac{\sum_{\mathrm{k}=1}^{\mathrm{n}} \mathrm{RMS}_{\mathrm{k}}^2}{\mathrm{n}}}$$

Where:

- $\mathrm{RMS_k}$: The RMS distance of the $k$-th residue as defined above.
- $n$: The total number of residues being compared.

**Log Probability (LogPr):**

The logPr metric quantifies the statistical likelihood of observing the angular alignment between two structures compared to a random environment. It uses the differences in backbone torsion angles ($\phi$ and $\psi$) between a reference and a fragment to compute a log-transformed probability [5].

$$\text{logPr} = \frac{1}{n} \sum_{k=1}^{n} \log_{10} \left( \frac{1}{180^\circ} \cdot \Delta\phi_k \cdot \frac{1}{180^\circ} \cdot \Delta\psi_k \right),$$

where:

- $n$: Total number of residues being compared.

- $\Delta\phi_k$: Absolute difference between the $\phi$ angles of the reference and fragment at residue $k$, with a small constant ($1e{-}10$) added to avoid zero.

- $\Delta\psi_k$: Absolute difference between the $\psi$ angles of the reference and fragment at residue $k$, with a small constant ($1e{-}10$) added to avoid zero.

#### 3 Number of Instance for Each Fragment

| Fragment Number | Count |
|---|---|
| 1 | 18 |
| 2 | 14 |
| 3 | 9 |
| 4 | 2 |
| 5 | 6 |
| 6 | 7 |
| 7 | 8 |
| 8 | 10 |
| 9 | 4 |
| 10 | 7 |
| 11 | 1 |
| 12 | 10 |
| 13 | 1 |
| 14 | 4 |
| 15 | 8 |
| 16 | 6 |
| 17 | 8 |
| 18 | 6 |
| 19 | 4 |
| 20 | 3 |
| 21 | 3 |
| 22 | 2 |
| 23 | 7 |
| 24 | 5 |
| 25 | 4 |
| 26 | 4 |
| 27 | 1 |
| 28 | 12 |
| 29 | 1 |
| 30 | 2 |
| 31 | 2 |
| 32 | 7 |
| 33 | 5 |
| 34 | 5 |
| 35 | 3 |
| 36 | 6 |
| 37 | 6 |
| 38 | 3 |
| 39 | 2 |
| 40 | 3 |

Table S1: Number of instances for each fragment in our fibrary.

### 4 Number of Proteins for each Function

| Function | Count |
|---|---|
| DNA+RNA | 10 |
| RNA | 10 |
| RNA+ATP+METAL | 10 |
| RNA+GTP | 10 |
| RNA+ATP | 10 |
| DNA+RNA+ATP | 9 |
| ATP+GTP+METAL | 10 |
| RNA+GTP+METAL | 10 |
| RNA+ATP+GTP | 10 |
| GTP | 11 |
| ATP+METAL | 10 |
| RNA+METAL | 9 |
| METAL | 10 |
| ATP | 9 |
| DNA+ATP+METAL | 10 |
| DNA+GTP | 6 |
| DNA | 8 |
| DNA+ATP+GTP | 5 |
| GTP+METAL | 10 |
| ATP+GTP | 9 |
| DNA+METAL | 10 |
| DNA+RNA+METAL | 9 |
| DNA+ATP | 10 |

Table S2: Number of Structures for each Function in the Protein Function Dataset (PFD).

### 5 Fragment Detection Accuracy

In this section, we assess the performance of our fragment detection algorithm by comparing predicted fragments against the ground truth library of verified instances derived from Alva et al. [1] from a held out test set. We evaluated both sequence-based (BLOSUM, Sequence Identity) and angle-based (RMS, RamRMSD, LogPr) distance metrics, as well as combinations thereof. The analysis focuses on determining the optimal metric combination via F1 scores and selecting the most robust distance threshold using Receiver Operating Characteristic (ROC) analysis.

##### 5.1 F1 Score

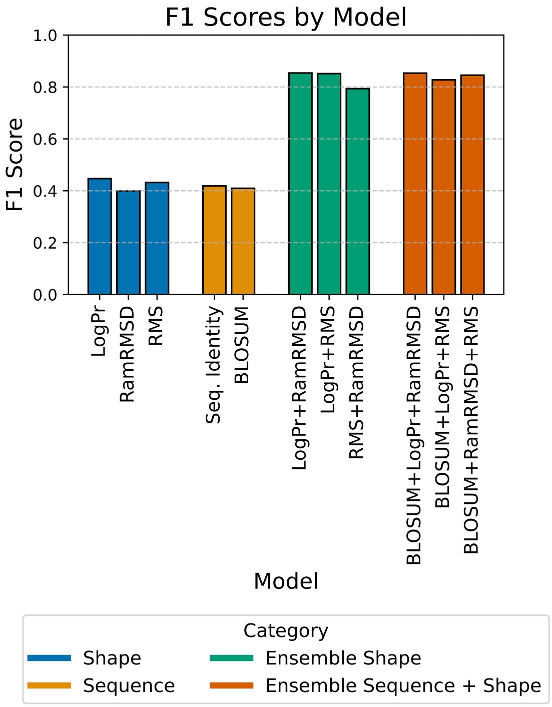


Figure S1: Comparison of F1 scores for different fragment detection models using various distance metrics. The figure illustrates the performance of individual sequence-based and angle-based metrics, as well as combined models.

##### 5.2 ROC Threshold

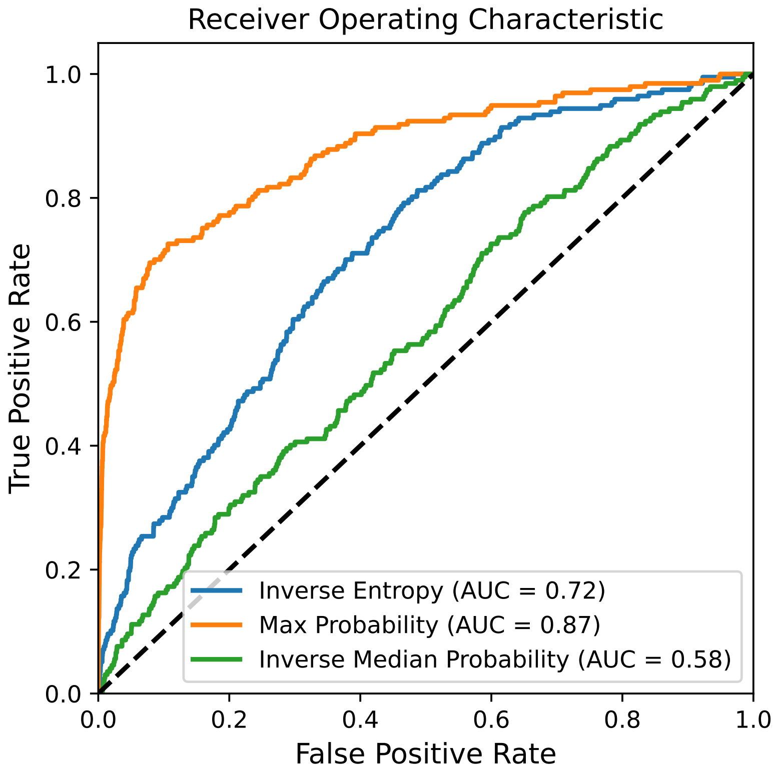


Figure S2: Receiver Operating Characteristic (ROC) curve for fragment detection. The ROC curve illustrates the true positive rate (sensitivity) versus the false positive rate (1 - specificity) for different classification thresholds. The area under the ROC curve (AUROC) quantifies the overall performance of the fragment detection algorithm, with higher values indicating better discrimination between true and false fragment classifications.

### 6 Distance Spearman Correlations

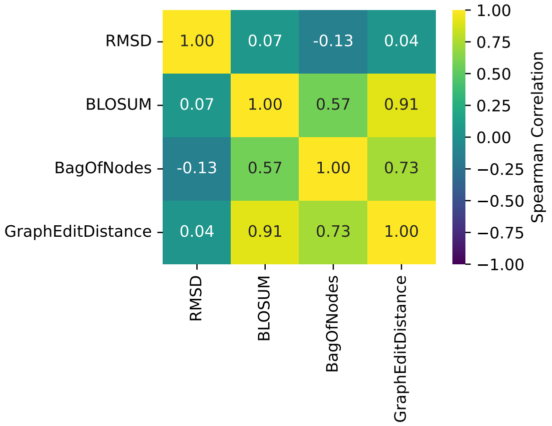


Figure S3: Spearman correlation matrix between different protein similarity metrics. The figure shows pairwise correlations between BagOfNodes (fragment set representation), Graph Edit Distance (GED, fragment graph representation), BLOSUM (sequence similarity), and RMSD (structural similarity). Spearman correlation values indicate the rank-based relationships between these metrics, capturing how similarity scores compare across different representations.

### 7 Functional Clustering

To evaluate the functional organization captured by our fragment-based representation, we performed unsupervised clustering on the embedding space. We benchmarked our fragment-based representations Graph Edit Distance and BagOfNodes, against standard structural (RMSD) and sequence-based (BLOSUM62) representations. For each metric, we computed pairwise distance matrices and projected the data into a 2-dimensional manifold using Principal Coordinate Analysis (PCoA), t-SNE, and UMAP. We then applied Gaussian Mixture Models (GMM) and K-Means clustering to these embeddings. The alignment between the resulting clusters and ground-truth functional labels was quantified using Adjusted Rand Index (ARI), V-Measure, and F1 scores, while the preservation of geometric topology was assessed via Trustworthiness and Spearman correlation (Table S3).

| Clustering Method | Dimensionality Reduction | Distance | ARI | V-Measure | Silhouette | Trustworthiness | Distance Spearman | F1 |
|---|---|---|---|---|---|---|---|---|
| GMM | PCoA | BagOfNodes | 0.0050 | 0.3455 | 0.8227 | 0.9991 | 0.9998 | 0.1660 |
| GMM | PCoA | BLOSUM | 0.0027 | 0.2933 | 0.0248 | 0.9923 | 0.9966 | 0.1640 |
| GMM | PCoA | GraphEditDistance | 0.0458 | 0.3832 | 0.0766 | 0.9915 | 0.9829 | 0.1985 |
| GMM | PCoA | RMSD | 0.0357 | 0.3863 | -0.0334 | 0.9598 | 0.8647 | 0.1957 |
| GMM | t-SNE | BagOfNodes | 0.0004 | 0.3431 | 0.8163 | 0.9945 | 0.9371 | 0.1485 |
| GMM | t-SNE | BLOSUM | 0.0043 | 0.3484 | -0.0004 | 0.8724 | 0.6607 | 0.1468 |
| GMM | t-SNE | GraphEditDistance | 0.0078 | 0.3499 | 0.1595 | 0.9787 | 0.8032 | 0.1632 |
| GMM | t-SNE | RMSD | 0.0370 | 0.3827 | 0.0006 | 0.6558 | 0.1752 | 0.1891 |
| GMM | UMAP | BagOfNodes | -0.0081 | 0.3283 | 0.6886 | 0.9466 | 0.5405 | 0.1478 |
| GMM | UMAP | BLOSUM | 0.0030 | 0.3452 | -0.0248 | 0.8717 | 0.6229 | 0.1392 |
| GMM | UMAP | GraphEditDistance | 0.0085 | 0.3545 | 0.1522 | 0.9707 | 0.5987 | 0.1656 |
| GMM | UMAP | RMSD | 0.0237 | 0.3741 | -0.0087 | 0.6681 | 0.1804 | 0.2038 |
| K-Means | PCoA | BagOfNodes | 0.0071 | 0.3486 | 0.8261 | 0.9991 | 0.9998 | 0.1562 |
| K-Means | PCoA | BLOSUM | 0.0020 | 0.2788 | 0.0238 | 0.9923 | 0.9966 | 0.1540 |
| K-Means | PCoA | GraphEditDistance | 0.0363 | 0.3621 | 0.1009 | 0.9915 | 0.9829 | 0.1868 |
| K-Means | PCoA | RMSD | 0.0486 | 0.3990 | -0.0249 | 0.9598 | 0.8647 | 0.2470 |
| K-Means | t-SNE | BagOfNodes | 0.0021 | 0.3459 | 0.8327 | 0.9945 | 0.9371 | 0.1635 |
| K-Means | t-SNE | BLOSUM | -0.0032 | 0.3375 | 0.0006 | 0.8724 | 0.6607 | 0.1670 |
| K-Means | t-SNE | GraphEditDistance | 0.0079 | 0.3536 | 0.1558 | 0.9787 | 0.8032 | 0.1782 |
| K-Means | t-SNE | RMSD | 0.0356 | 0.3859 | 0.0111 | 0.6558 | 0.1752 | 0.2248 |
| K-Means | UMAP | BagOfNodes | -0.0061 | 0.3278 | 0.6735 | 0.9466 | 0.5405 | 0.1406 |
| K-Means | UMAP | BLOSUM | 0.0023 | 0.3481 | -0.0126 | 0.8717 | 0.6229 | 0.1516 |
| K-Means | UMAP | GraphEditDistance | 0.0024 | 0.3462 | 0.1520 | 0.9707 | 0.5987 | 0.1490 |
| K-Means | UMAP | RMSD | 0.0240 | 0.3745 | 0.0032 | 0.6681 | 0.1804 | 0.1782 |

Table S3: Clustering performance across different methods, distance metrics, and dimensionality reduction techniques. The table reports Adjusted Rand Index (ARI), V-Measure, Silhouette score, Trustworthiness, Spearman correlation with original distances, and F1 score for Gaussian Mixture Models (GMM) and K-Means clustering. Each method is evaluated using four different distance metrics—BagOfNodes, BLOSUM, Graph Edit Distance (GED), and RMSD—applied to embeddings obtained via Principal Coordinate Analysis (PCoA), t-SNE, and UMAP.

#### 8 Search Performance

We use the following metrics:

- **Normalized Discounted Cumulative Gain (NDCG)**: evaluates the quality of ranking by comparing the actual ranking with an ideal one, emphasizing higher relevance at the top of the list. It calculates a weighted relevance score for ranked proteins, where the highest possible score (ideal) is compared to the actual retrieval. NDCG accounts for multi-functional proteins by assigning partial relevance to those containing the query function.

- **Area Under the Receiver Operating Characteristic (AUROC)**: measures how effectively the method ranks proteins sharing the same function higher than those with different functions. For each protein in the dataset, we identify relevant proteins (i.e., those matching the query's function) and compute the AUROC by comparing the distances between proteins, with closer distances indicating higher relevance. Distances are converted into scores, and the AUROC quantifies the likelihood that a relevant protein is ranked above a non-relevant one. This metric is aggregated across all proteins and functional categories in the dataset to assess overall retrieval success.

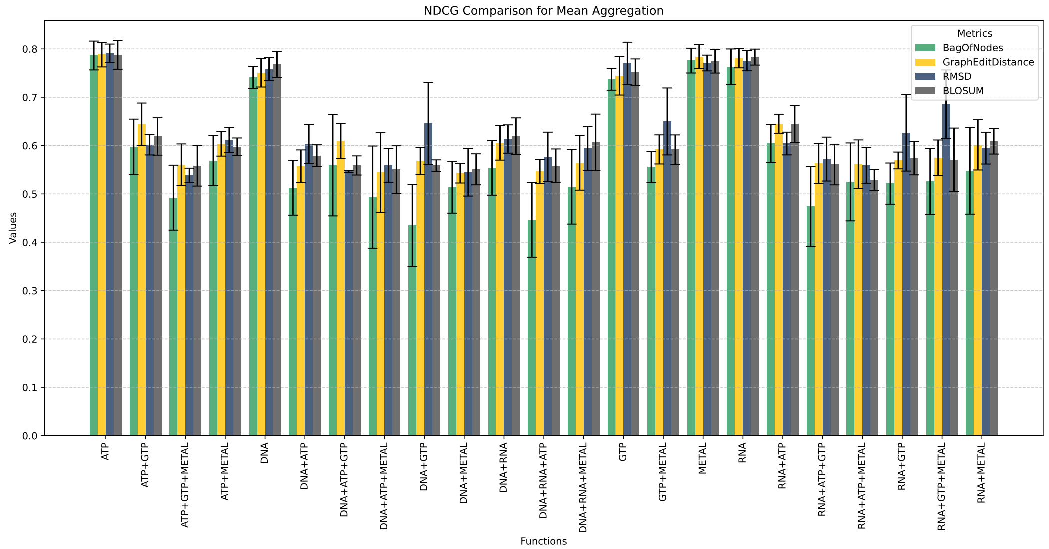


Figure S4: Normalized Discounted Cumulative Gain (NDCG) scores for protein similarity search across different distance metrics. Higher values indicate that functionally relevant proteins are ranked higher in the search results.

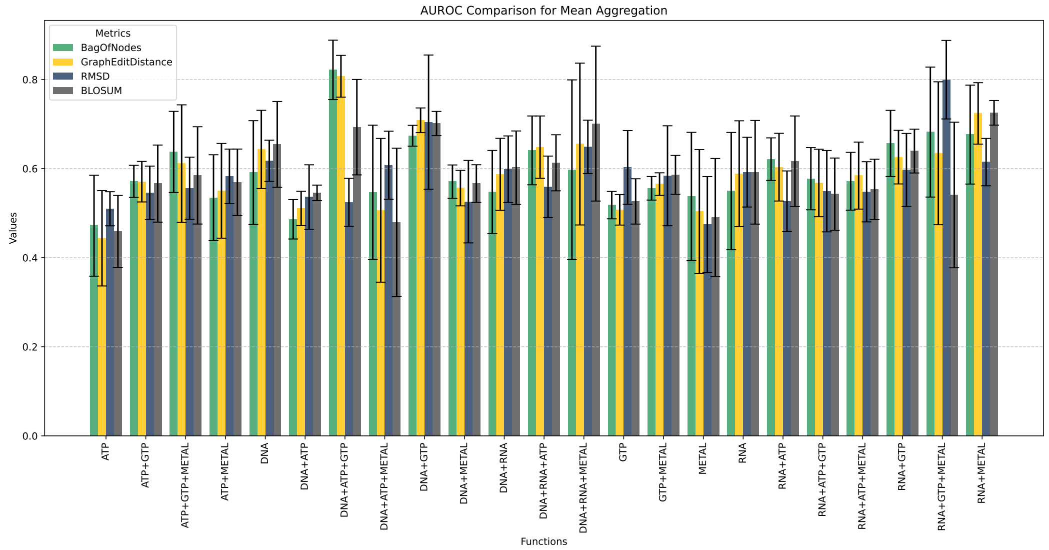


Figure S5: Area Under the Receiver Operating Characteristic Curve (AUROC) scores for protein similarity search across different distance metrics. Higher values indicate better separation between relevant and non-relevant search results.

### 9 Physico-chemical Properties in Fragments

Fragment coverage quantifies the proportion of a protein structure that is classified to a known fragment in our representation. This analysis evaluates fragment coverage across different protein folds and resolutions using the PDBench dataset[2]. Additionally, we investigate how fragment coverage correlates with key structural and chemical properties, such as hydrogen bonding, solvent accessibility, charge distribution, polarity, and secondary structure content.

To assess these relationships, we compare coverage across different fold classes and resolution ranges, and we analyze whether fragment regions exhibit distinct physicochemical characteristics compared to non-fragment regions.

Fragment coverage for binary properties (e.g., charge, polarity, secondary structure) is calculated as the fraction of all residues possessing a specific property that fall within fragment regions. For continuous properties like relative solvent accessibility (RSA), we calculate the proportion of the total cumulative RSA value contributed by fragment residues compared to the whole protein.

To evaluate structural independence, we decomposed hydrogen bonds using the Kabsch-Sander algorithm into two categories: **intra-fragment bonds** (where both donor and acceptor reside within the same fragment instance) and **inter-fragment bonds** (connecting different fragments).

Finally, to normalize for varying fragment density across proteins, we computed the **Coverage Ratio** defined as:

$$\text{Ratio} = \frac{\text{Coverage}_{\text{prop}}}{\text{Coverage}_{\text{fold}}}$$

A ratio greater than 1.0 indicates that the property is enriched within fragment regions, while a ratio less than 1.0 indicates depletion relative to the rest of the backbone.

##### 9.1 Fragment Coverage By Fold and Resolution

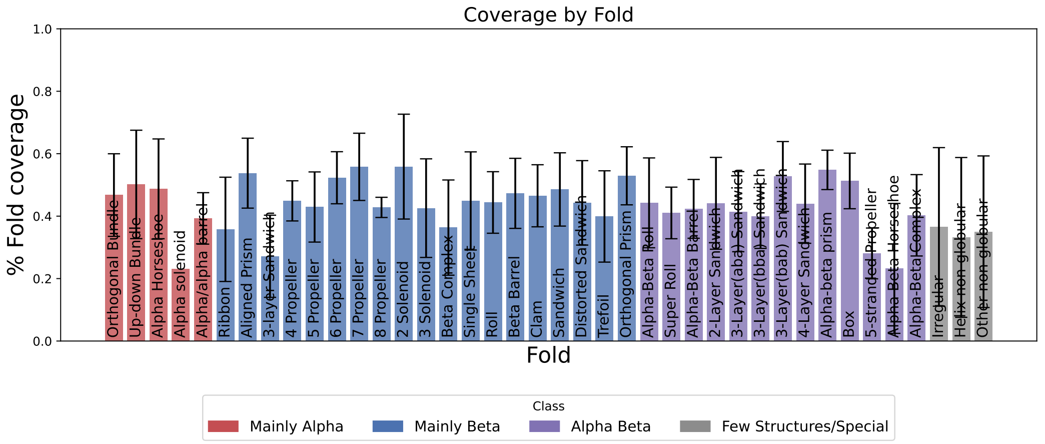


Figure S6: Coverage of Fragments Across Folds

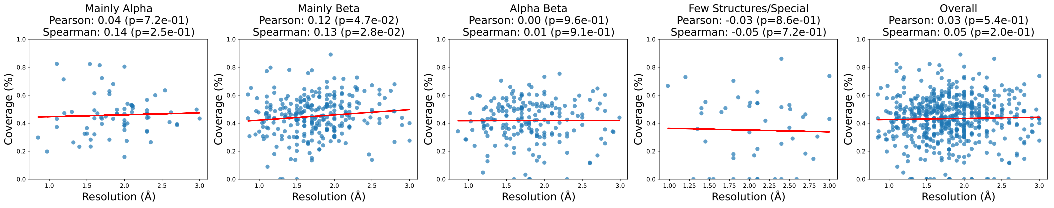


Figure S7: Coverage of Fragments Across Resolutions

#### 9.2 H-bond Between Fragments Coverage

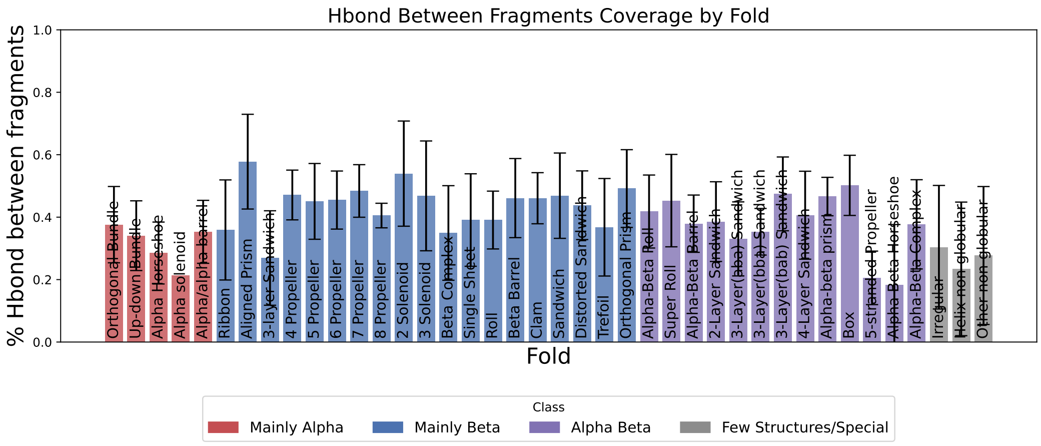


Figure S8: H-bond Between Fragments Coverage by Fold.

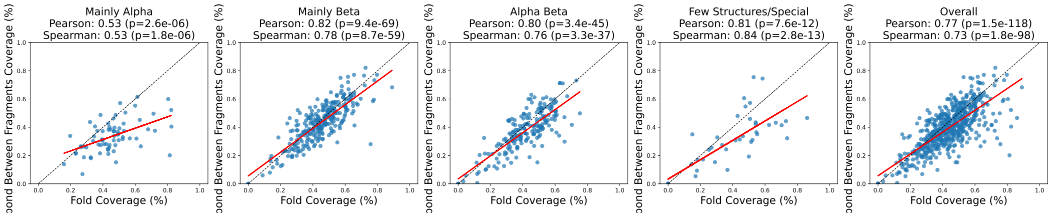


Figure S9: H-bond Between Fragments vs. Fold Coverage.

#### 9.3 H-bond Within Fragments Coverage

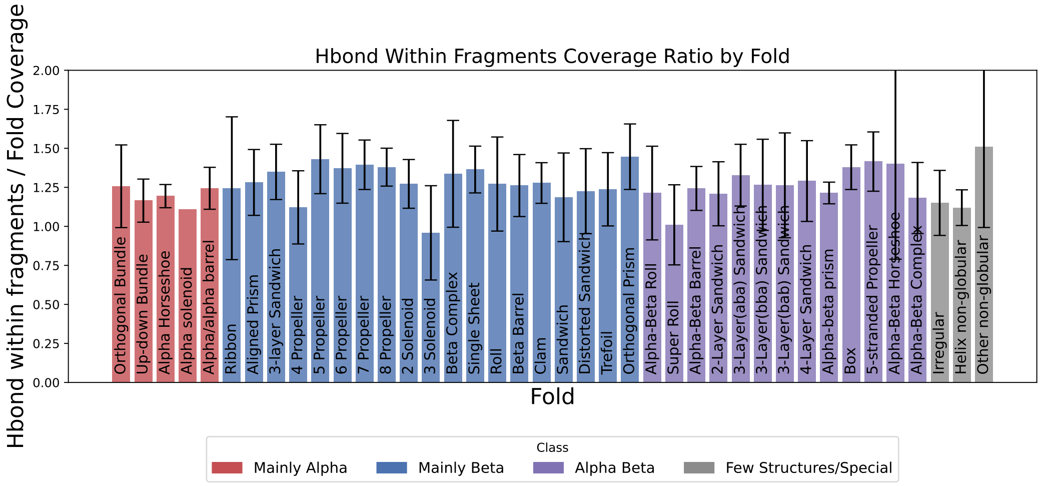


Figure S10: H-bond Within Fragments Coverage Ratio by Fold.

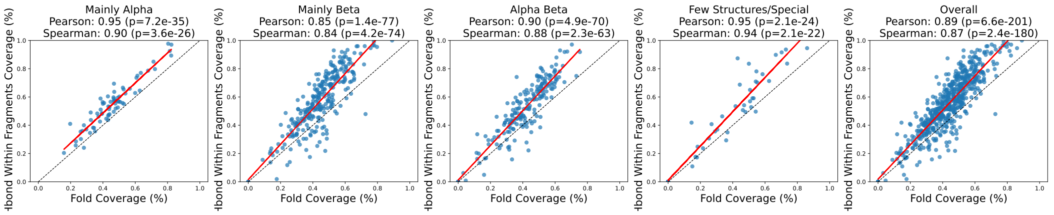


Figure S11: H-bond Within Fragments vs. Fold Coverage.

#### 9.4 Accessibility Coverage

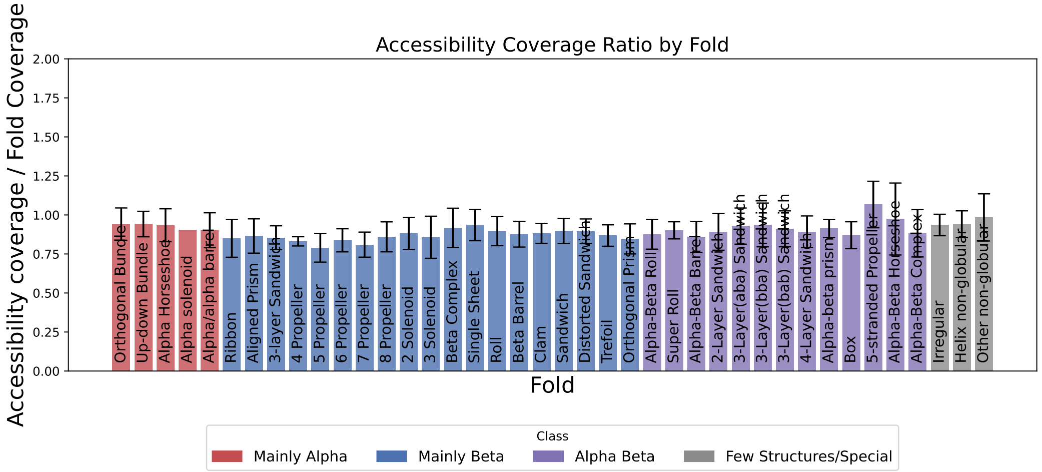


Figure S12: Accessibility Coverage Ratio by Fold.

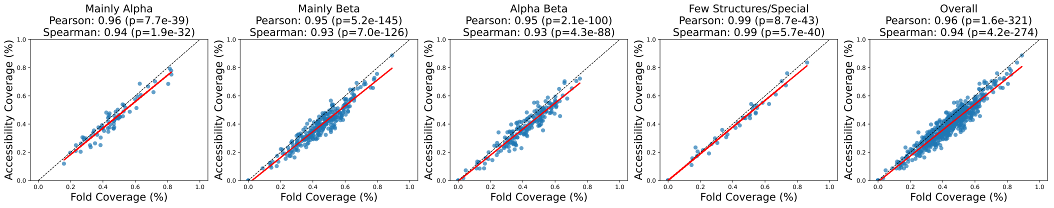


Figure S13: Accessibility vs. Fold Coverage.

#### 9.5 Charge Coverage

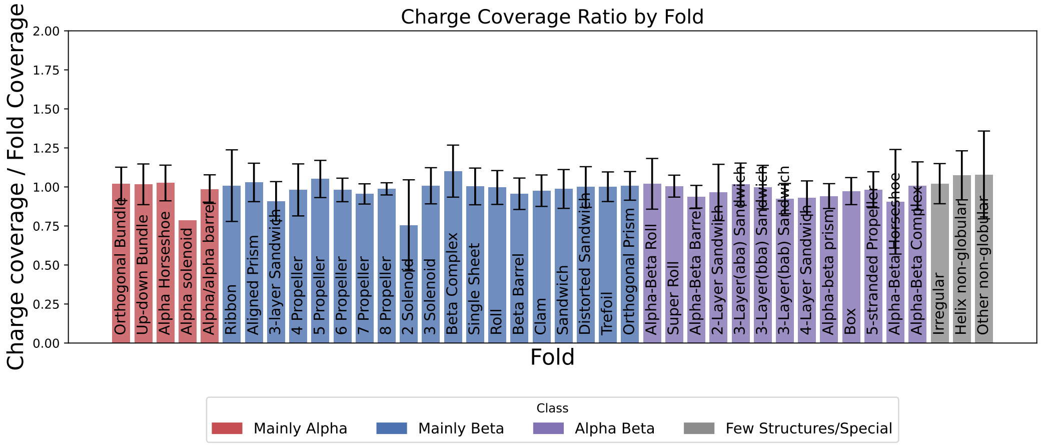


Figure S14: Charge Coverage Ratio by Fold.

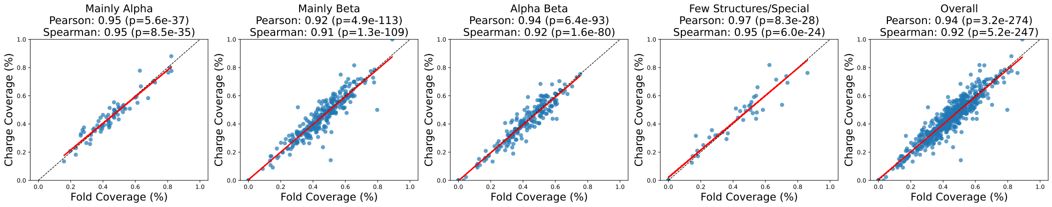


Figure S15: Charge vs. Fold Coverage.

#### 9.6 Polarity Coverage

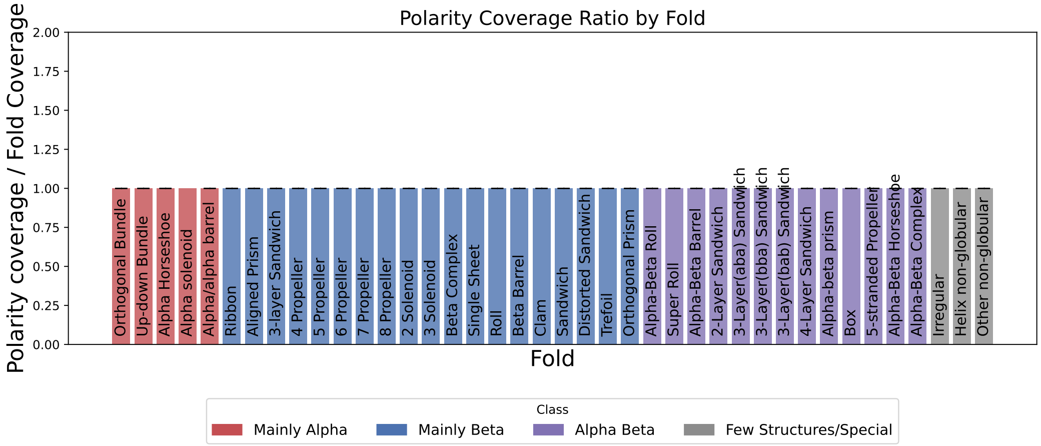


Figure S16: Polarity Coverage Ratio by Fold.

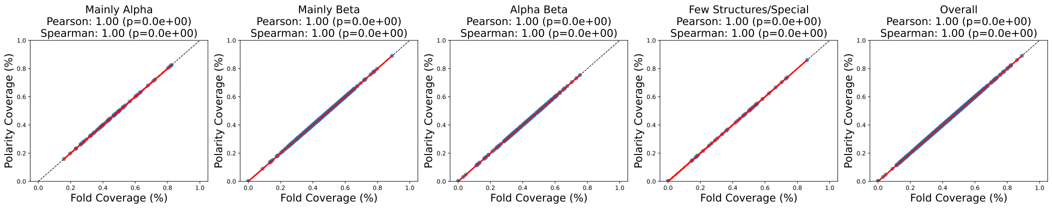


Figure S17: Polarity vs. Fold Coverage.

#### 9.7 Secondary Structure Coverage

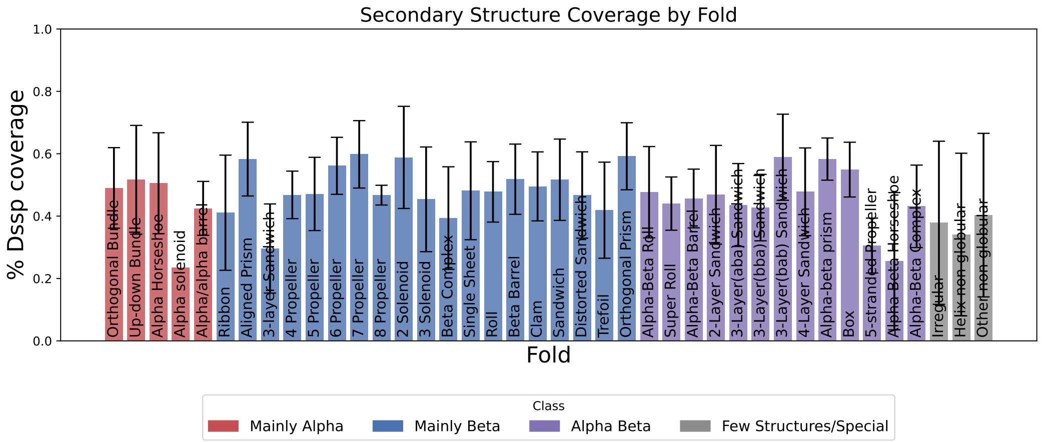


Figure S18: Secondary Structure Ratio by Fold.

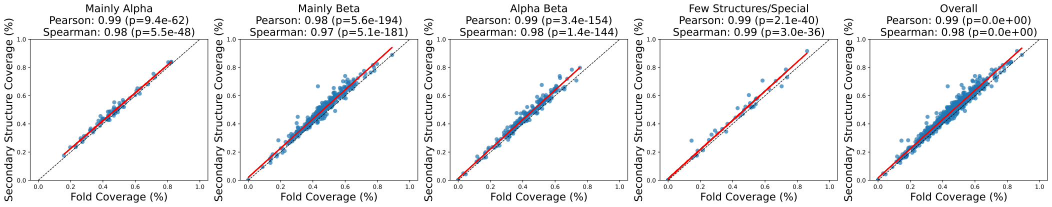


Figure S19: Secondary Structure vs. Fold Coverage.

### 10 Design Success Rate

In this section, we evaluate the generative capacity of our fragment representation. We used the detected evolutionary fragments as fixed geometric constraints for the generative model RFDiffusion, tasking it with inpainting the missing backbone regions to create complete protein structures. The functional fidelity of these de novo designs was assessed by querying the generated backbones against the PDB and SwissProt using FoldSeek with TM-Align mode. A design was considered successful if its top 10 structural matches shared the same Gene Ontology (GO) function as the original template protein, measuring whether the newly diffused structures with fragment templates were sufficient to encode functional geometry.

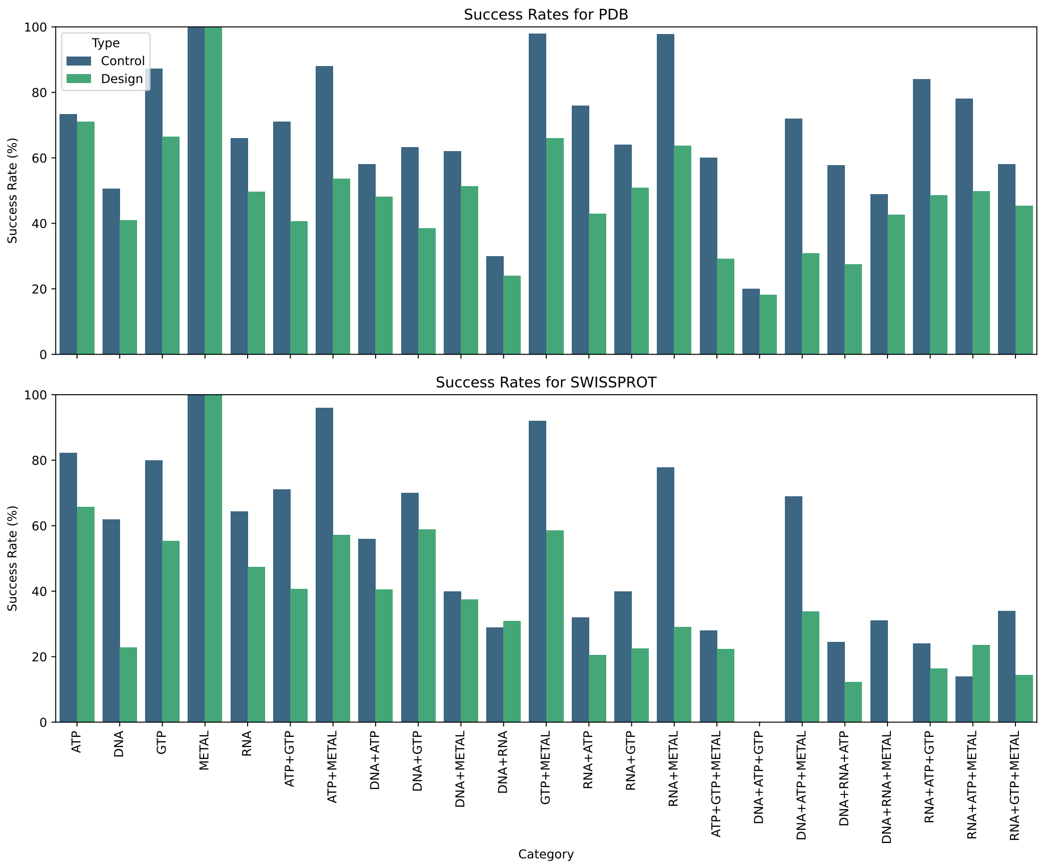


Figure S20: Success rates for fragment-constrained backbone generation. Each protein was used as a template for generating five backbone designs, guided by detected functional fragments. The success rate is defined as the proportion of generated backbones whose top 10 structural matches in FoldSeek share the same Gene Ontology (GO) function as the original protein.

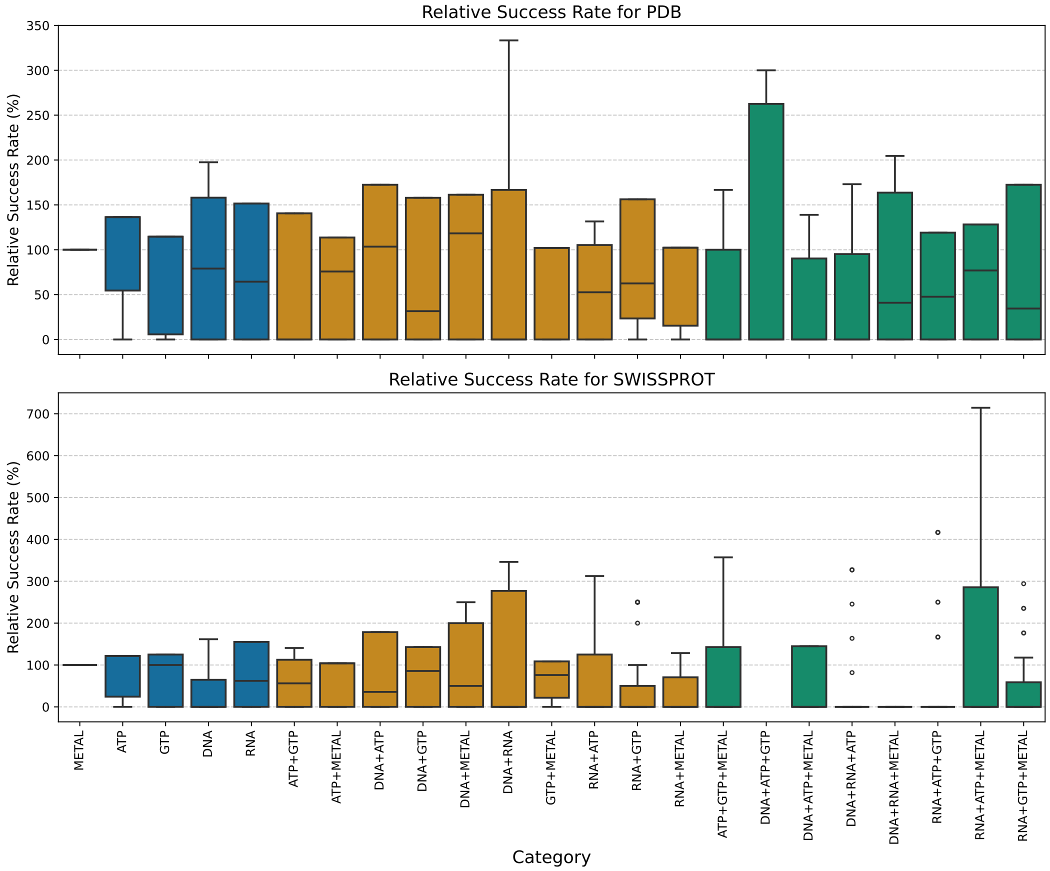


Figure S21: Boxplot of relative success rates for fragment-constrained backbone generation. The relative success rate compares the recovery of functionally similar proteins between generated backbones and their original templates. Values greater than 100 indicate improved functional recovery over the original structure, while values below 100 suggest reduced functional specificity in the generated designs.

images/relative_success_rates.pdf

Figure S22: Relative recovery rates for fragment-constrained backbone generation. The fragments from each protein in the Protein Function Dataset (PFD) were used to generate a template for RFDiffusion. The recovery rate is defined as the fraction of generated backbones whose top 10 structural matches in FoldSeek share the exact Gene Ontology (GO) function of the original protein. The relative recovery rate compares this to the recovery rate of the original protein backbone.

##### 10.1 Validation against Naive Control

To validate that the functional recovery observed above is driven by the specific evolutionary information encoded in our library, rather than simply by geometric constraints, we performed a control experiment using "naive" fragments. These were generated by extracting random, non-overlapping sequence windows from the native backbone, matching the exact length and residue count of the evolutionary fragments used in the main experiment.

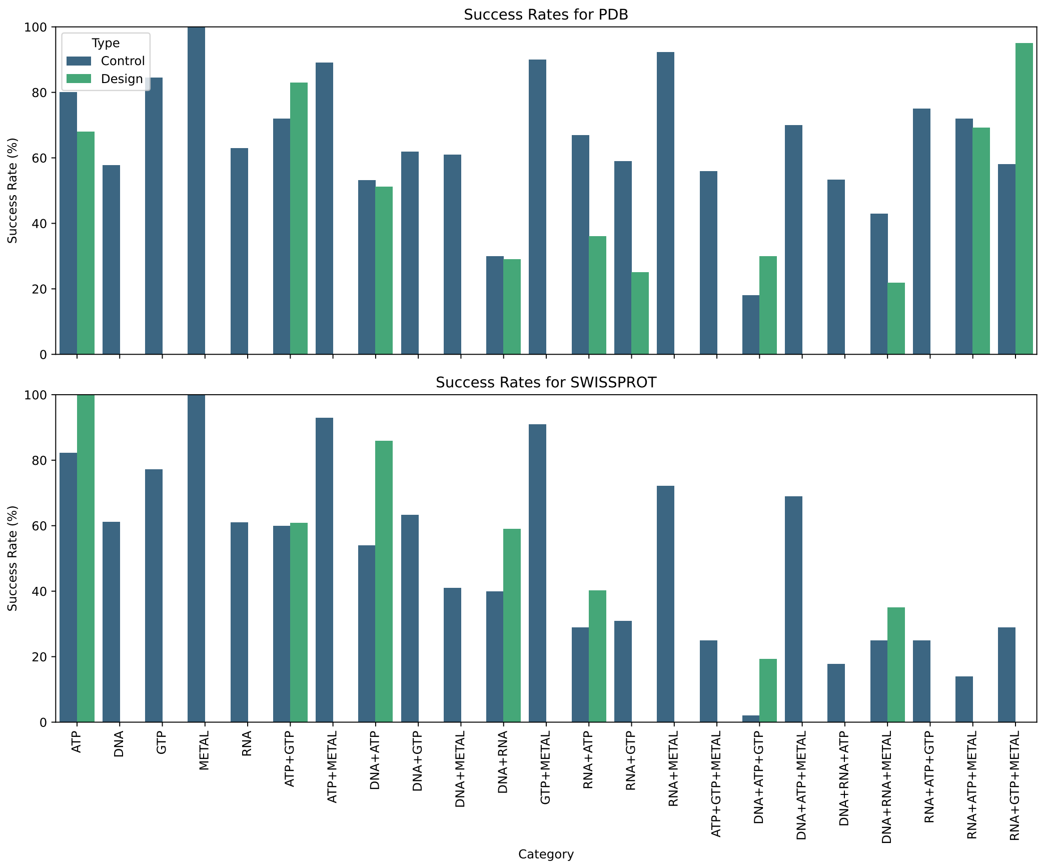


Figure S23: Success rates for naive control backbone generation. Each protein was used as a template for generating five backbone designs, guided by *random* backbone constraints (naive fragments).

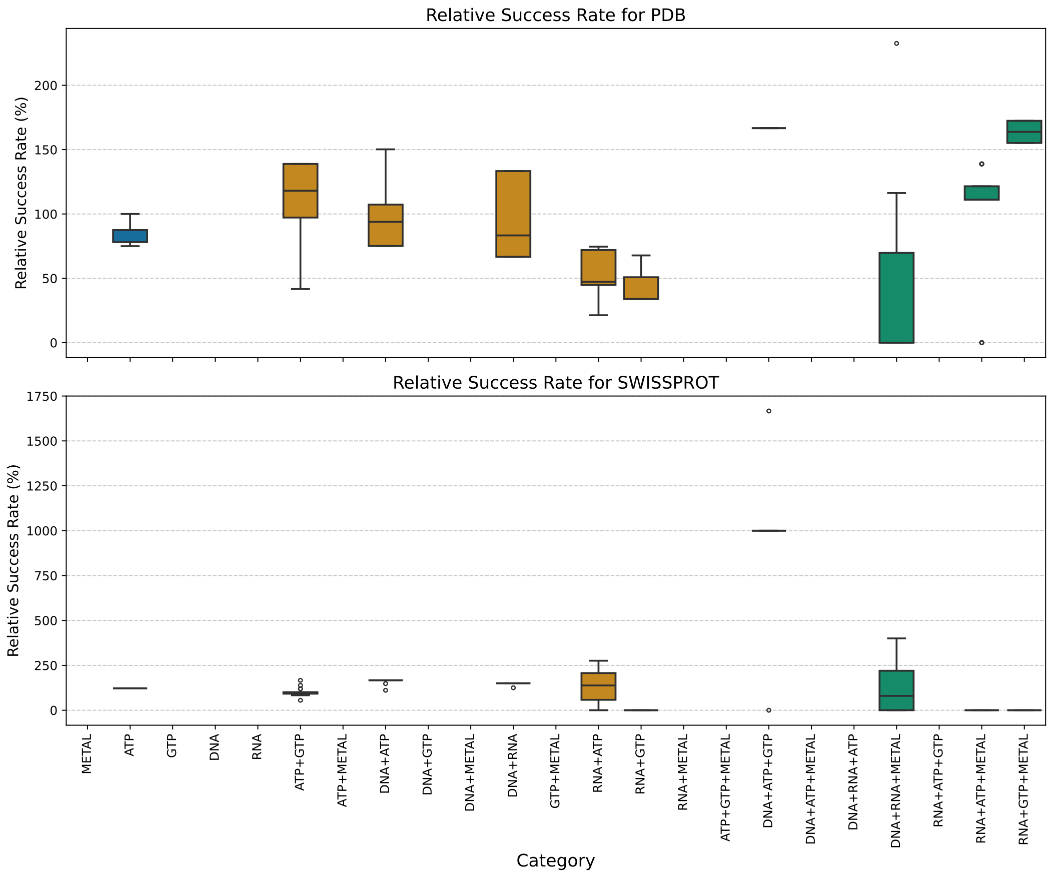


Figure S24: Boxplot of relative success rates for naive control.

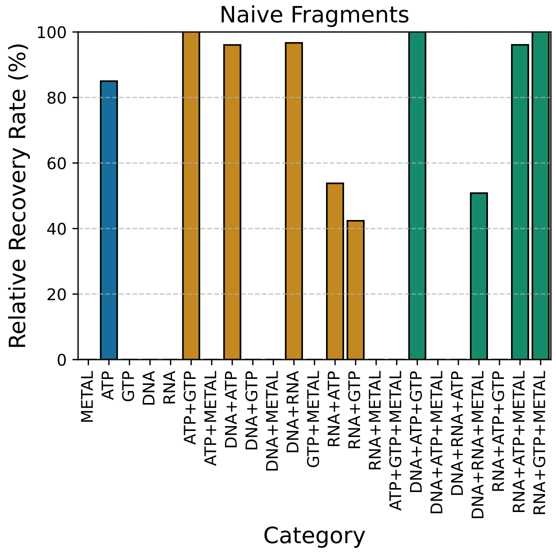


Figure S25: Relative recovery rates for naive control backbone generation. Random fragment constraints from each protein were used to guide RFDiffusion. While robust folds (e.g., ATP-binding) show high recovery, geometry-sensitive functions such as Metal, DNA, and RNA binding show near-zero relative recovery (compare to Fig. S22).

#### 11 Data Point Distributions of Fragments and traditional methods

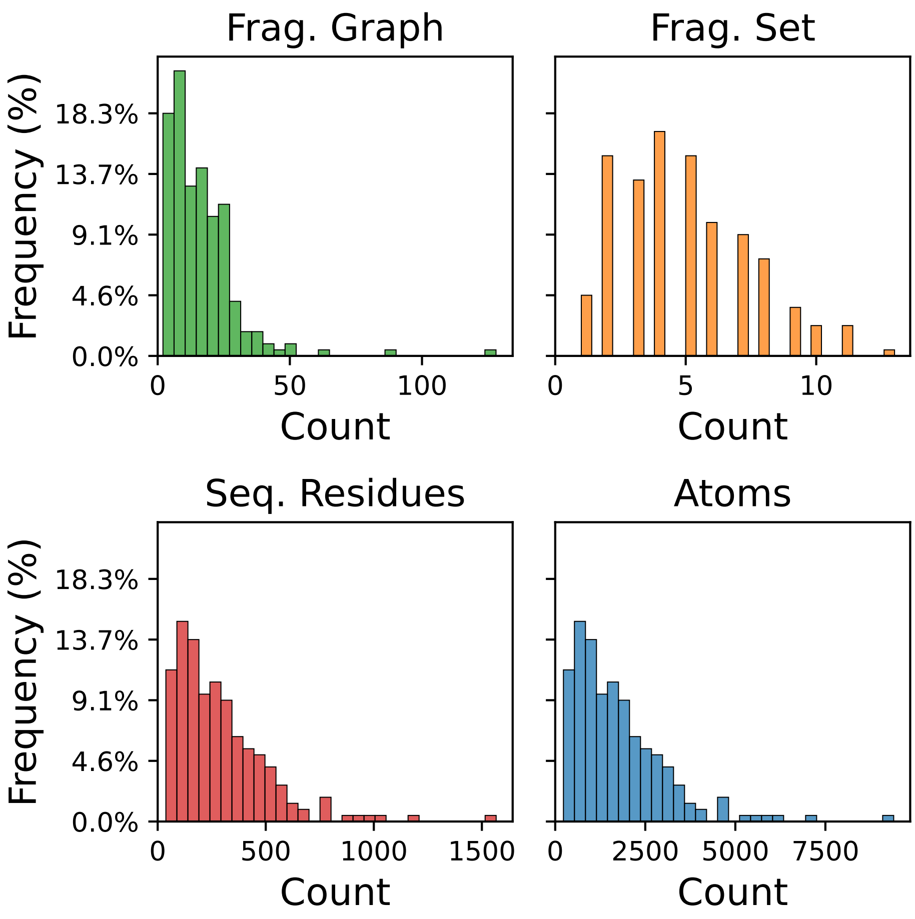


Figure S26: Distribution of the number of data points required to represent proteins using different methods: Fragment Nodes (Fragment Graphs), Fragment Sets, Residues (Sequence-based representation), and Atoms (Shape-based representation). The figure illustrates how different levels of abstraction affect the number of data points required for each representation, highlighting variations in data complexity across methods.

#### 12 Qualitative Analysis

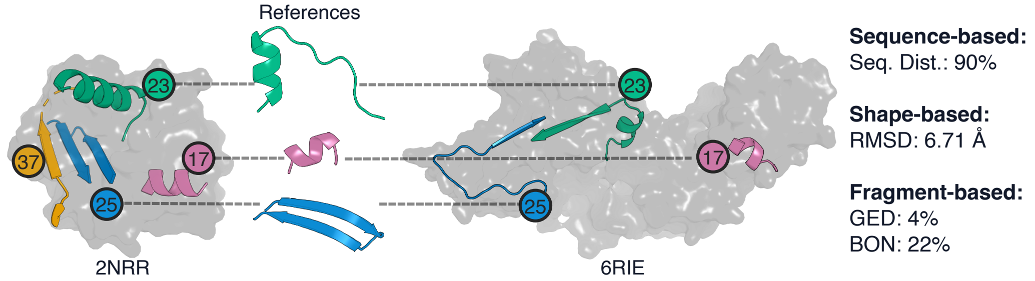


Figure S27: Comparison of two structurally distinct DNA-binding proteins using sequence-, shape-, and fragment- based methods. The UvrABC system protein C, involved in DNA repair (PDB: 2NRR), is shown on the left, while a viral DNA-dependent RNA polymerase (PDB: 6RIE) is on the right. Despite significant differences in sequence and overall structure, both proteins share common functional fragments (17, 23, and 25, highlighted in color). Traditional sequence- and structure-based distances indicate high divergence between these proteins. In contrast, fragment-based metrics show a relatively low distance, suggesting a potentially shared functional role.
